# Porcine Parvovirus VP1/VP2 on a Time Series Epitope Mapping: exploring the effects of high hydrostatic pressure on the immune recognition of antigens

**DOI:** 10.1101/330589

**Authors:** Ancelmo Rabelo de Souza, Marriam Yamin, Danielle Gava, Janice Reis Ciacci Zanella, Maria Sílvia Viccari Gatti, Carlos Francisco Sampaio Bonafe, Daniel Ferreira de Lima Neto

**Affiliations:** Departamento de Bioquímica e Biologia Tecidual e, Universidade Estadual de Campinas (UNICAMP), Rua Monteiro Lobato, 255, Cidade Universitária Zeferino Vaz, 13083-862, Campinas, SP, Brazil; Departamento de Genética, Evolução e Bioagentes, Instituto de Biologia, Universidade Estadual de Campinas (UNICAMP), Rua Monteiro Lobato, 255, Cidade Universitária Zeferino Vaz, 13083-862, Campinas, SP, Brazil; Embrapa Suínos e Aves, Laboratório de Virologia de Suínos, 89715-899, Concórdia, SC, Brazil

**Keywords:** epitope mapping, epitope prediction, molecular dynamics, porcine parvovirus, spot synthesis

## Abstract

Porcine parvovirus (PPV) is a DNA virus that causes reproductive failure in gilts and sows, resulting in embryonic and fetal losses worldwide. Epitope mapping of PPV is important for developing new vaccines. In this study, we used spot synthesis analysis for epitope mapping of the capsid proteins of PPV (NADL-2 strain) and correlated the findings with predictive data from immunoinformatics. The virus was exposed to three conditions prior to inoculation in pigs: native (untreated), high hydrostatic pressure (350 MPa for 1 h) at room temperature and high hydrostatic pressure (350 MPa for 1h) at −18 °C, compared with a commercial vaccine produced using inactivated PPV. The screening of serum samples detected 44 positive spots corresponding to 20 antigenic sites. Each type of inoculated antigen elicited a distinct epitope set. *In silico* prediction located linear and discontinuous epitopes in B cells that coincided with several epitopes detected in spot synthesis of sera from pigs that received different preparations of inoculum. The approach used here provided important information on the antibody/antigen interactions required to improve B cell responses to PPV and may be useful in developing novel strategies for producing new vaccines.

**Abbreviations:** 3D, three dimensional; HHP, high hydrostatic pressure; ID, identification; IEDB, Immune Epitope Database; ORFs, open reading frames; p.i., post-infection; PPV, porcine parvovirus; RMSD, root-mean-square deviation of atomic positions; R(g), radius of gyration. RMSF, root-mean-square fluctuation; SK6, Swine kidney cell; SPF, specific pathogen free; TCID^50^/mL, Median Tissue Culture Infectious Dose; HI, Hemagglutination Inhibition.

## 1. Introduction

Porcine parvovirus (PPV), or Ungulate Protoparvovirus 1 as proposed by Cotmore et al. ^1^, is a 25-nm diameter, non-enveloped icosahedral virus that contains ∼5 kb of negative sense, single-strand DNA (ssDNA) comprising two large open reading frames (ORFs) in its genome. ORF1 codes for the nonstructural proteins NS1, NS2 and NS3, and ORF2 codes for the structural proteins VP1, VP2 and VP3 ^2^. VP1 and VP2 capsid proteins are the result of an alternative splicing of the same gene and VP3 itself is formed by proteolytic cleavage of VP2. These structural proteins are responsible for the immunogenic properties of PPV ^3^. PPV infects pregnant gilts and sows, causing reproductive failure characterized by embryonic and fetal death, mummification and stillbirths, with delayed return to oestrus ^4^. The resulting reduction in reproductive capacity can significantly decrease pork production ^5^.

PPV is prevalent in the pig population and highly stable in the environment, which make it difficult to establish and keep breeding populations free of the virus. Based on this, it is important to maintain herd immunity against PPV ^4^. The most widely used approach for maintaining immunity is regular vaccination of breeding females. Two main strains of PPV have been identified, based on pathogenicity: NADL-2 (non-pathogenic) and Kresse (pathogenic), although sequence analysis of recent isolates suggests active evolution of PPV ^6^. The vaccines commercially available since the 1980s are based on a chemically-inactivated NADL-2 strain.

The development of vaccines is important for controlling diseases such as reproductive failure caused by PPV. In this context, essential considerations should include viral diversity, protective immunity and population coverage as key factors, all of which pose special challenges to this ever-expanding field ^7,8^. One well-known approach in vaccine production is the use of inactivated viral preparations ^9,10^. In recent years, the use of high hydrostatic pressure (HHP) has become an increasingly popular non-thermal method of inactivation ^11-13^. This approach has been successfully used to inactivate several viruses and represents a promising alternative to vaccine development ^14^. Several studies have demonstrated that pressure can result in virus inactivation while preserving immunogenic properties. In addition, HHP is a technology free of chemicals that is safe and capable of inducing strong humoral and cellular immune responses. In this study, we used a combination of *in silico* and *in vivo* approaches to examine the immune responses to different HHP-PPV formulations that were then compared with a commercial vaccine produced using chemically-inactivated virus.

## 2. Material and methods

### 2.1 Virus preparation

The NADL-2 strain of PPV was cultured in SK6 cells as previously described^15^. Supernatants of cell cultures were separated and purified by CaCl_2_ precipitation followed by gradient ultracentrifugation, as previously described ^16^. Fractions with hemagglutinating activity were dialyzed for 16 h at 4 °C against a Tris-EDTA buffer, pH 8.0 ^17^. The virus was titrated in SK-6 cells by serial dilutions in 96 well-plates and the cells were then screened for a cytopathic effect at 72 h post-infection. The test was considered positive when cytopathic effects were observed in >75% of the cells. Samples containing 1×10^4.5^ TCID_50_/mL were aliquoted and subjected to HHP. Vaccine aliquots and viral preparations were normalized to avoid an antigen-concentration effect ^18^.

Briefly, virus aliquots were subjected to two treatments: one at 300 MPa for 1 h at 25 °C and another at 300 MPa at −18 °C, both immersed in 0.1 M Tris-HCl, pH 7.4. The equipment used to treat the samples consisted of a pressure generator device (model HP ISS) coupled to a pressure cell, as previously described ^19^. The freezing point (i.e., −19 °C) for these conditions was determined to be lower in previous studies by our group ^20^.

### 2.2 Pig immunization and sample collection

This experimental study was done at the Embrapa (Brazilian Agricultural Research Corporation) Swine and Poultry Station, located in the southern Brazilian state of Santa Catarina. All the experimental protocols involving animals were approved by the Institutional Animal Care and Use Committee (Embrapa, protocol n° 003/2009).

Ten 21-day-old specific-pathogen-free (SPF) male pigs from a terminal cross of Landrace and Large White lineages were divided into five groups of two pigs each:(N) (native PPV), (P) (PPV treated with 350 MPa at 25 °C), (P-18) (PPV treated with 350 MPa at −18 °C), (V) (commercial inactivated PPV vaccine, Farrow sure B, Pfizer) and (NC) (negative control). Pigs were vaccinated intramuscularly on day 0 (D0), D14, D28 and D38 according to each HHP protocol, as previously described ^50^. At necropsy following euthanasia (D72), samples of lung, heart, liver, spleen, kidney and mesenteric lymph node were collected and stored at −70 °C until analyzed by nested-PCR. Pigs in the NC group received only saline solution. On D51, all pigs were challenged intranasally with a reference strain of PPV NADL-2, as previously described^21^. Serum samples were collected prior to each vaccination or challenge (D0, D14, D28, D38 and D51), as well as on D58, D65 and D72. On the last day of the experiment, a post-mortem examination was done and samples of lung, heart, liver, spleen, kidney and mesenteric lymph node were collected and stored at −70°C until analyzed by nested-PCR.

### 2.3 Detection of DNA and anti-PPV antibodies

Viral DNA was extracted from pig tissues using DNeasy Blood & Tissue kits (Qiagen) and subjected to nested-PCR for PPV detection, as previously described ^22^. Pig serum samples were tested for anti-PPV antibodies by an HI test ^23^.

### 2.4 Spot synthesis

The peptides were synthesized based on the amino acid sequence of VP1 from PPV NADL-2 strain (GenBank, accession no. NC 001718) with 12 overlapping peptides and an offset of four peptides. The membrane used for spot synthesis was prepared with Fmoc (9-flurorenylmethoxycarbonyl) chemistry ^24,25^ on a PEG-derived cellulose membrane with the addition of a C-terminal anchor of β-Ala residues and hexadecapeptides at the N-terminal. All serum samples from D0, D28, D58 and D72 (representing, respectively, the basal serum condition prior to any intervention, the antibody response against the vaccine, the challenge with the reference strain and 21 days after the challenge) were screened for antibodies. An anti-swine IgG-Fc secondary antibody conjugated with alkaline phosphatase (Jackson ImmunoResearch Europe Ltd.) was used to assess epitope recognition in the peptide arrays and downstream procedures were done as previously described ^25^. The membrane was subsequently scanned at 1200 d.p.i. with a Print Scan Copier (HP Photosmart) and the software package “Totallab Quant-Array Analysis” was used to quantify the spots based on the intensity of the reaction relative to the background intensity.

### 2.5 Linear and conformational epitope prediction

The complete primary sequence of the capsid protein of the reference strain was obtained from the UniProt Knowledge Base (UniProtKB) database (accession no. P18546) and used for downstream applications. Linear TCD4^+^ and TCD8^+^ epitope prediction was done using the tools of the Immune Epitope Database (IEDB) with the sequence of the reference strain used in the experiments as input. The physicochemical properties of amino acids were considered and assessed using the Karplus and Schulz flexibility scale ^26^. The antigenicity of the protein was inferred with the Kolaskar and Tongaonkar scale, a semi-empirical method to predict antigenicity from the physicochemical properties of amino acids in protein sequences ^27,28^. The hydrophobic properties of the sequence were predicted with the Parker hydrophilicity scale ^29^.

### 2.6 Molecular dynamics

The crystallized structure of the VP2 protein (PDB code 1k3v) was downloaded from the Protein Data Bank. Molecular dynamics simulations were done with the GROMACS software (v. 5.1.3) ^30^ to evaluate putative structural changes caused by HHP (350 MPa) at 27 °C and −18 °C relative to the control conditions of atmospheric pressure at 27 °C. The system consisted of 184,609 atoms with 58,720 water molecules and 8,447 atoms representing the 1k3v solvated protein coordinates after energy minimization; eight chloride ions were added to stabilize the system. For the calculations, a cubic simulation box with a periodic boundary cell of 80 × 80 × 80 nm was assumed. The CHARMM22 force field was applied for 1k3v and chloride ions and the TIP3P water model were selected for the nvt and npt equilibration protocols. The electrostatic interactions were calculated by using the particle mesh Ewald (PME) method with a cut-off distance of 13.0 Å for the van der Waals interactions and the electrostatic interactions in the real space of the PME method. The temperatures and pressures were selected using the Nosé-Hoover Langevin piston pressure control based on the Nosé-Hoover thermostat and the Langevin piston method. To simulate the experimental conditions the equilibration systems were normalized to 300 K at atmospheric pressure followed by variations in the nvt and npt protocols to achieve the desired HHP conditions, i.e., 350 MPa at 27 °C and −18 °C. Production runs for each condition were done for 20 ns and the outputs analyzed for structural parameters such as root mean square deviations (RMSD), root mean square fluctuations (RMSF), radius of gyration (Rg), secondary structure fluctuations and also for the complete system, such as pressure, temperature and energy variations. Plots were generated with the Grace software package (http://plasma-gate.weizmann.ac.il/Grace/) and trajectory analyses were done using the VMD package ^31^ as well as the built-in analysis packages present in GROMACS ^30^.

## 3. Results

### 3.1 PPV DNA and antibody detection

Table 1 shows the detection of PPV DNA in different tissues. Groups N and NC were PPV-positive in spleen, liver, lymph node and lung. Group V was positive only in lymph node and kidney. Both HHP groups (P and P-18) were positive in spleen and liver, whereas lung positivity was identified in group P and in lymph node in group P-18. No viral DNA was detected in heart samples of any group.

**Table 1.**
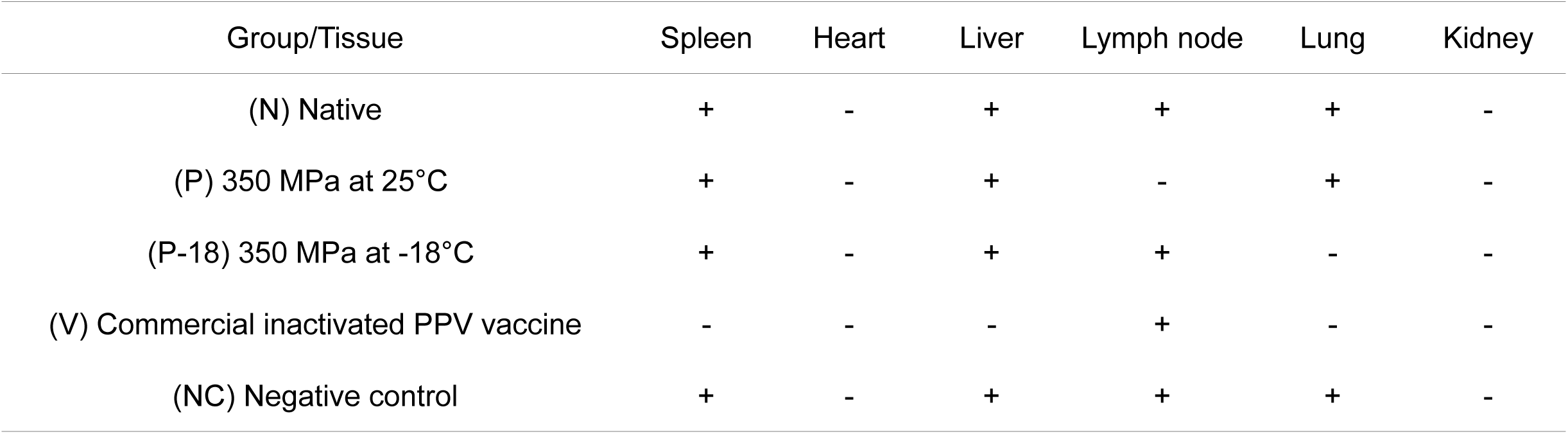

No pigs presented anti-PPV antibodies at the beginning of the experiment (D0). By D14 after the first vaccination, low titer anti-PPV antibodies started to be detected. Pigs inoculated with HHP (groups P and P-18) had higher antibody titers than the pigs vaccinated with reference strain NADL-2. As expected, by D72 after the PPV challenge, all pigs seroconverted, with titers varying from 1024 to 2048.

### 3.2 Epitope mapping

A schematic representation of the membrane is depicted in Figure 1A, with the positive spots highlighted in Figure 1B and C, portraying the detailed conditions for each positive result (different HHP treatments and serum sampling). None of the serum samples collected at D0 bound any polyclonal antibodies. Negative control pigs (group NC) showed positivity only after PPV inoculation (D58), which reflected sites usually activated when natural PPV infection occurs (sites 13 and 16 that correspond to amino acid residues 493-512 and 585-608, respectively) (Figure 1C).

**Fig. 1.**
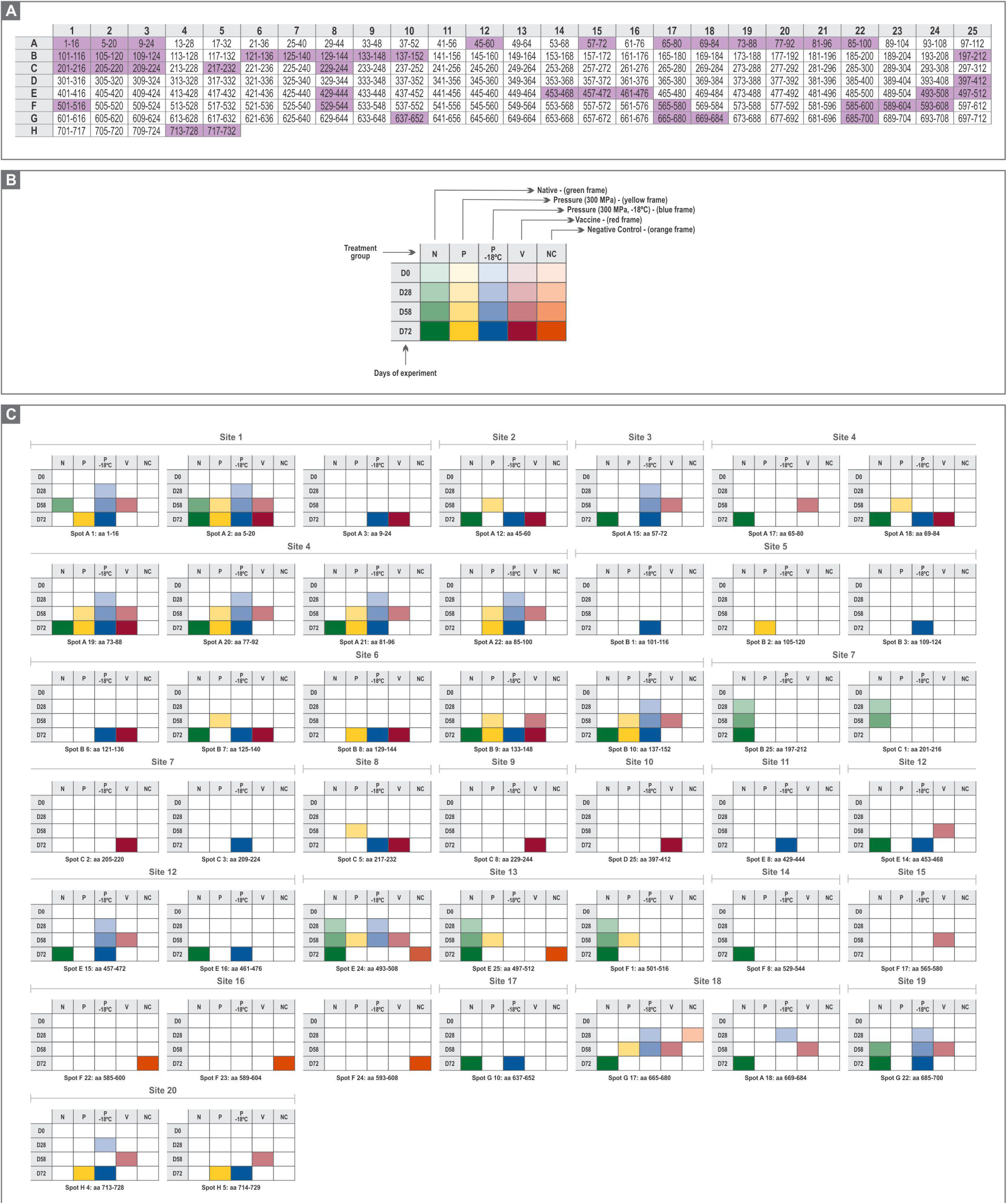
Epitope mapping results for the different conditions of pig inoculations. (A) Membrane spot array with each hexadecapeptide showing the VP1 amino acid position in each cell. The highlighted spots (green, n=44) correspond to positivity in at least one of the experimental conditions. (B) An explanatory framework for panel C.(C) Detailed data of the spots highlighted in A, showing the corresponding positivity for the experimental conditions and serum samples (3, 6 and/or 8). Consecutive positive spots were denominated “sites” (n=20).

The data for each experimental condition and time point are shown as pairs of graphs in Figure 2. The spot intensity is shown and was considered positive in Figure 2A, C, E, G and I; the corresponding location was mapped onto the crystal structure of PPV VP2, with color coding according to the time point (Figure 2B, D, F, H and J). There was a predominance of red and yellow regions in several conditions, indicating that the initial positivity in a given region frequently did not persist after challenge. In addition, spots that were positive after challenge (sample 8) were not preceded by positive epitope mapping in initial inoculations.

**Fig. 2.**
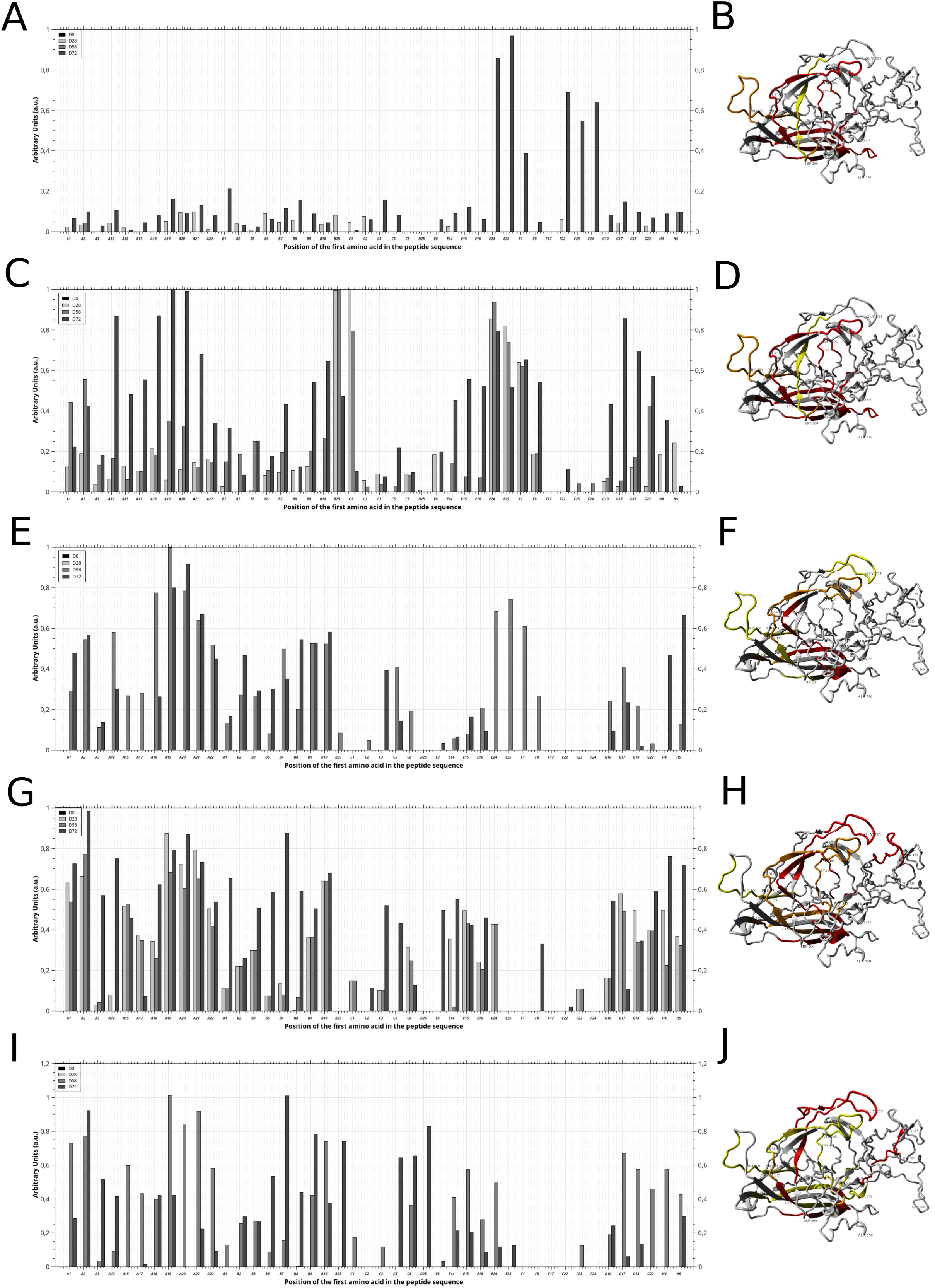
Intensity of the colorimetric output of the antibody-bound alkaline phosphatase reactions (panels A, C, E, G and I) and structural representation of each mapped region on the crystal structure of VP2 for each condition (panels B, D, F, H and J). The signal intensity of the positive spots shown in Figure 1C, based on serum samples 3 and 6, and the post-challenge sample (8), was used to construct epitope maps of PPV. The data were normalized relative to the maximum intensity for each experimental condition. The threshold was considered 40% of maximum intensity. The models on the right show the crystallographic structure of VP2 protein (PDB entry 1k3v) with positivity indicated as yellow for sample 3 and/or 6, red for sample 8, and orange for positivity in both 3 and/or 6 and 8.

Pigs from group N showed no significant reaction on D28, except for sites 7 and 13 that correspond to amino acid residues 197-216 and 493-516, respectively. When samples from D58 (representing the immune response after PPV challenge) were analyzed, additional positivity occurred at sites 1 (aa 1-20) and 19 (aa 685-700). When antibodies obtained after the challenge with active virus were mapped against the membrane (D72), numerous additional epitope sites were detected, namely, 2 (aa 45-60), 3 (aa 57-72), 4 (aa 65-96), 6 (aa 125-140; 133-152), 12 (aa 453-476), 14 (aa 529-544), 17 (aa 637-652) and 18 (665-684). Interestingly the region initially detected at site 7 (aa 201-216) on D28 and D58 was not detected in the last sample (D72).

For group V, immunoreactivity was observed only on D58. In this case, PPV antibodies were detected at more sites than with the native form of the virus. Sites 3 (aa 57-72), 4 (aa 65-80; 73-100), 6 (aa 133-152), 12 (aa 453-472), 15 (aa 565-580),18 (aa 665-684) and 20 (aa 713-729) had bound antibodies, in contrast to group N for which no signal was found in the same regions at D58. Epitope mapping after the viral challenge (D72) showed higher agreement with the map of group N, but distinct regions were also found for sites 1 (aa 9-24), 6 (aa 121-136; 129-144), 7 (aa 205-220), 8 (aa 217-232), 9 (aa 229-244) and 10 (aa 397-412).

In general, a higher number of sites was induced and recognized in group P than in group N, but this number was lower than in group V. Sites 1, 4, 6, 13 and 18 present in group P were also found in samples 3 and 6 of group V that had been inoculated intramuscularly. Sites 1 and 13 were also detected in group N. At the last sample collected (D72), the native form showed more antigenic sites compared with groups P and V (Table 1S and Figure 1C). Site 1 showed the highest frequency of positive spots at D28, D58 and D72 in groups N, P and V (Table 1S and Figure 1C).

In group P-18, there was an increase in the number of positive spots (Figure 1C). This situation was similar to group N, but the mapped response was higher when P-18 was compared to groups P, V and NC. Group P-18 pigs showed positive antibody responses earlier than the other treatments and included sites 1, 3, 4, 6, 12,18, 19 and 20 (Figure 1C). This was the only in which sites 5 (aa 101-116; 109-124)7 (aa 209-224) and 11 (aa 429-444) were activated at D72.

### 3.3 Bioinformatic analysis

A combination of immunoinformatic tools and molecular dynamics simulations was used to understand the molecular recognition of the VP1-VP2 capsid proteins based on the epitope mapping results described above. By using the reference sequence of NADL-2 strain, *in silico* predictions identified sites with a high probability of being antigens. Since this method does not evaluate tertiary structures and is based on a sliding window to produce propensity scores, we identified six regions out of the 24 predictions that overlapped with the results for group N.

Based on properties such as solvent-accessible surface area (SASA) and hydropathy predictions (Supplementary Figure 1 (SF-1)) these regions were found to be exposed and hydrophilic, thus providing basic information for the experimental detection of such regions as epitopes in group N. Considering the conformational epitope predictions and conservative settings used, few regions returned positive conformational epitope predictions, but balancing for sensitivity and specificity the algorithm was able to detect putative epitopes on the 1k3v reference structure (public data), albeit on distinct regions when compared with our epitope mapping of the untreated virus.

Bioinformatics analysis with GROMACS allowed simulation of the behavior of VP2 under normal conditions and under the conditions of groups P and P-18. A 20 ns molecular dynamics simulation allowed comparison of the radius of gyration (Rg), an indicator of protein compactness, with the group N simulation and revealed an overall decrease of 0.42% in Rg for group P *vs.* group N (2.85 nm *vs.* 2.838 nm) and a 0.87% decrease for the P-18 simulation compared to group N (2.85 nm *vs.* 2.826 nm) (SF1), suggesting that the simulation produced a more constrained protein. The total energy of the HHP simulation increased 0.83% against group N (−2.385 × 10^6^ cal *versus* −2.405 × 10^6^ cal) whereas that for group P-18 increased 9.85% (−2.385 × 10^6^ cal *versus* −2.635 × 10^6^ cal) (SF1). Conversely, a marked decrease in total volume was recorded during the group P and P-18 simulations compared with group N, which fluctuated around 1835 nm^3^ *vs.* 1680 nm^3^ (P) and 1655 nm^3^ (P-18), suggesting that the simulation was successful in creating the desired pressure and temperature for the experiment (SF-1, SF-2).

The RMS fluctuation after the production run revealed the baseline values for the native protein atom fluctuations in accordance with the predicted flexible regions of the protein and provided a basis for comparison with the group P and P-18 simulations. As a result, the P simulation narrowed the amplitude variation per atom, thereby collapsing the flexible regions and limiting the overall variation to 0.01 nm. Furthermore, the P-18 simulation produced a more rigid configuration with no fluctuations >0.05 nm on all atoms, in agreement with the other parameters (total volume, total energy and Rg). Interestingly, the area per residue over the trajectory increased substantially in the group P and P-18 simulations (SF-2). Moreover, the number of contacts maintained between residues decreased substantially when groups P and P-18 were compared with control group N (SF-2).

## 4. Discussion

Epitope mapping strategies have been extensively applied to key pathogens and the results provided by this approach have important applications in public health, as well as in animal safety and welfare ^32^–^36^. There are ongoing efforts to identify the key epitopes on the VP1-VP2 capsid proteins of PPV through distinct strategies for mapping. A study in which pepscan was used to synthesize 24 different peptides directed towards antigenic sites detected nine such sites and posited that those found in the N-terminal region were neutralizing epitopes ^37^. Xie et al.^3^ confirmed these findings by using a monoclonal antibody (C4) against VP1 of PPV to screen a 12-mer phage peptide library. These authors identified a mimotope corresponding to the N-terminal immuno-dominant region that protected mice against the virus. In the present study, we mapped 20 “antigenic sites” corresponding to 44 positive spots based on a combination of experimental conditions that corresponded to vaccination/challenge settings. Distinct patterns of epitope positivity were observed, depending on the experimental conditions to which the antigen was subjected and on whether the vaccination or challenge phase was being assessed (Figure 2A-J).

To our knowledge, spot synthesis methodology has not previously been applied to PPV in this manner and the results obtained here provide a comprehensive landscape of the immunological profile of the VP1-VP2 complex by slicing the protein into overlapping oligopeptides bound to a nitrocellulose membrane. The spot results are quantifiable and provide satisfactory parallel comparisons with ELISA findings, such as for determining protein concentration ^38^; the methodology is reproducible and has been previously used for quality control ^39^.

By combining molecular dynamics with epitope mapping, it was possible to investigate the putative alterations that HHP may have induced in protein conformation and in the solvent-accessible surface area. The native form of the protein, used here as a positive control, provided the epitope landscape to which pigs responded at three sampling times. Initially, group N showed a strong antibody response for spots B25 (aa 197-212) and C1 (aa 201-216) at sampling times D28 and D58. These spots correspond to the N-terminal region of VP2, information that is also present in the crystal structure of VP2. Epitopes located in the VP2 capsid protein of PPV are important for generating neutralizing antibodies against this virus, with a high potential to activate B cells. These studies indicate that the N-terminal of VP2 is a strong candidate for B cell epitopes and should be included in vaccines ^40,41^. Antibodies that bound to a region composed of amino acids 497-516 were seen soon after the second inoculation and the titers (inferred from the intensity of the phosphatase reactions) were maintained throughout the experiment. The immunodominant regions in VP2 belong to the loops and other regions in the N- and C-terminals ^37,42,43^. As shown in Figure 2B, D, F, H and J, the loop regions marked by Pro^500^ and Val^175^ occurred in epitopes present in the five groups, with an alternation of these regions in three intervals of serum collection.

Although the capsid proteins were deemed to be under near-neutral selection, a few key amino acids were found to be variable (aa 215, 228, 383, 414, 419 and 436), with most of them located in surface loops ^43,44^. Zeeuw et al.^45^ have previously shown that the strain switch from avirulent (143a) to virulent (27a) might depend directly on mutations of VP2 residues 378 and 383. As expected, surface amino acids in the VP2 configuration are hydrophilic and contain most of the mutations identified by Ren et al. ^46^, who also found this protein to be under negative selection. Antibody responses against overlapping peptides of VP1 were not detected until late time points, precisely after the viral challenge, when amino acids 5 through 96 produced significant binding, thereby reinforcing previous findings regarding the N-terminal immunogenicity of PPV VP1 ^3^.

In contrast to these results, vaccine epitope mapping elicited marked responses against the N-terminal region of VP1 after D38, with moderate binding to the previously mentioned VP2 regions. HHP elicited responses like those of the vaccine in the VP1 N-terminal region, as well as for the VP2 variable regions mentioned before on the fourth inoculation, prior to the challenge with PPV. In our study, amino acid residues (45, 217-219 and 556) corresponding to the mapped epitopes included some of the sites reported by Streck et al. ^44^. The substitution of amino acid residues at the sites located mainly on the surface of PPV are important not only for understanding the evolutionary mechanism of adaptive response to the host, but also for identifying possible strategies to produce more effective vaccines against PPV.

HHP treatment of PPV at 25 °C and -18 °C did not suppress antigenicity but elicited the appearance of several other epitopes through the exposure of parts of the capsid. Although HHP did not directly affect the tertiary structure of the proteins, the exposure of different parts of the proteins, including hydrophobic regions, in response to these conditions could explain the appearance of new epitopes ^47^. A direct comparison of the secondary structures of the VP2 crystal structure obtained after a 20 ns molecular dynamics simulation of the experimental conditions also showed small alterations to the original secondary structure configuration, mostly from switches between flexible regions (the most frequent interconversions found were from coil to bend, turn to3-helix and alpha-helix to turn to 3-helix). Conversely, HPP and low temperature HPP produced more stable secondary structures compared to the simulation with no pressure or temperature constraints. HHP and low temperature HHP simulations displayed small fluctuations during the production run and only the N-terminal region of the VP2 1K3V structure showed changes in configuration (amino acids 40-90). Compared with the molecular dynamics of native VP2, the HHP simulation tended to produce fixed turns where interconversion between bends and turns were recorded (amino acids 45-55) whereas low temperature HHP fixed this region in a 3-helix configuration. The region from amino acids 75-85 fluctuated among α-helix/turn/3-helix configurations in the native simulation but the region was then fixed as an α-helical structure in the HHP run and varied between turns and bends in the low temperature HHP run.

Previous reports have identified important structural changes in globular proteins after exposure to HHP, with increased disordered structures and turns, as well as decreased β-sheets and α-helices ^13^. These findings suggest that the effect of pressure on kinetics arises from a larger positive activation volume for folding than for unfolding; this in turn leads to a significant slowing down of the folding rate with increasing pressure. For viral capsids under HHP, D’Andrea et al ^48^ noted that the treatment enhanced the viricidal effect on hepatitis A virus, a non-enveloped virus, indicating that the efficacy of treatment depends on capsid conformation. These authors suggested a correlation between mature and immature capsids, with the accessibility of the immunodominant site near the five-fold axis possibly explaining the susceptibility of the virus to inactivation by HPP ^48^.

Under HHP, a wide variety of experimental data indicates the formation of a small cavity in globular proteins, capable of accommodating water, and the protein interior is associated with elastic deformation, the contribution of which to free energy considerably exceeds the heat motion energy. This hypothesis has an important consequence for globular proteins based on the mechanical results of the process when related to the intramolecular dynamics of proteins, namely, that the mobility of ions from the solvent to the interior of a protein should depend on pressure, obeying its intrinsic forces at low pressure whilst enhancing them at high pressure ^49^.

Current methods of vaccine production make use of hazardous chemicals that may pose a risk to vaccine recipients. The efficacy of HHP for inactivating a variety of foodborne pathogenic microorganisms is well established, and some of these microorganisms have been demonstrated to retain immunogenic properties, a finding which suggests that HHP may have an application in vaccine development. Indeed, pressure can result in virus inactivation while preserving immunogenic properties^11,12,14,50^.

Viruses contain several components that can be susceptible to the effects of pressure. HHP has been a valuable tool for assessing viral structure-function relationships because the viral structure is highly dependent on protein-protein interactions. In the case of small icosahedral viruses, incremental increases in pressure produce a progressive decrease in the folding structure when moving from assembled capsids to ribonucleoprotein intermediates (in RNA viruses), free dissociated units (dimers and/or monomers) and denatured monomers. High pressure inactivates enveloped viruses by trapping their particles in a fusion-like intermediate state. The fusogenic state, which is characterized by a smaller viral volume, is the final conformation promoted by HHP, in contrast to the metastable native state, which is characterized by a larger volume. The combined effects of high pressure with other factors, such as low or subzero temperature, pH and agents in sub-denaturing conditions, such as urea at low concentration, have been a formidable tool for assessing the component’s structure, as well as pathogen inactivation ^14,51^. HHP allows the production of inactivated vaccines that are free of chemicals, safe and capable of inducing strong humoral and cellular immune responses.

Inactivated or attenuated strains of PPV for vaccine production may contribute to the control of this virus in pigs. In this context, epitope mapping of immunogenic proteins of PPV is crucial for assessing immune recognition of such viral preparations with a potential use in vaccine preparation, as shown here for the immunization of pigs. In the present case, the combination of HHP and epitope mapping provided us with a broader, more refined perspective of the epitope landscape stimulated by each preparation. This information could potentially be used to track HHP-vaccinated pig herds as a first scenario, or, conversely, be used as a combined alternative to cover the immune response against the virus. Taken together, our findings provide detailed insight into the immune system of pigs when the animals are stimulated with the same molecular configuration of an antigen, the structure of which can be altered by pressure and temperature. These data also improve our understanding of site-directed antibody production over time and of the putative immunological dynamics of antibody selection before and after a challenge with antigen.

## Acknowledgements

The authors thank Embrapa (Concórdia, SC, Brazil) for providing the piglets and the molecular data and Stephen Hyslop for editing the English of the manuscript. This study was supported by Fundação de Amparo à Pesquisa do Estado de São Paulo (FAPESP, grant no. 2008/09835-0) and Coordenação de Aperfeiçoamento de Pessoal de Nível Superior (CAPES, grant no. grant no. 2012/1143817).

---

**Supplementary figure 1.** – Molecular dynamics production run results: First line: results for 20 ns of production run of the 1k3v negative control. A – Volume, B – Temperature, C – Pressure. Second line: results for 20 ns of production run of the 1k3v simulated under pressure and low temperature. D – Volume, E – Temperature, F – Pressure. Third line: results for 20 ns of production run of the 1k3v simulated under pressure: G – Volume, H – Temperature, I – Pressure.

**Supplementary figure 2.** – Molecular dynamics production run results: First line: results for 20 ns of production run of the 1k3v negative control. A – Solvent Accessible Surface Area per residue, B – RMSF per residue, C – number of contacts. Second line: results for 20 ns of production run of the 1k3v simulated under pressure and low temperature. D – Solvent Accessible Surface Area per residue, E – RMSF per residue, F – number of contacts. Third line: results for 20 ns of production run of the 1k3v simulated under pressure: G – Solvent Accessible Surface Area per residue, H – RMSF per residue, I – number of contacts.

